# Dynamics of venom composition across a complex life cycle

**DOI:** 10.1101/159889

**Authors:** Yaara Y. Columbus-Shenkar, Maria Y. Sachkova, Arie Fridrich, Vengamanaidu Modepalli, Kartik Sunagar, Yehu Moran

**Author notes:** These authors contributed equally to this work (Yaara Y. Columbus-Shenkar, Maria Y. Sachkova).

## Abstract

Little is known about venom in young developmental stages of animals. The appearance of stinging cells in very early life stages of the sea anemone *Nematostella vectensis* suggests that toxins and venom are synthesized already in eggs, embryos and larvae of this species. Here we harness transcriptomic and biochemical tools as well as transgenesis to study venom production dynamics in *Nematostella*. We find that the venom composition and arsenal of toxin-producing cells change dramatically between developmental stages of this species. These findings might be explained by the vastly different ecology of the larva and adult polyp as sea anemones develop from a miniature non-feeding mobile planula to a much larger sessile polyp that predates on other animals. Further, the results suggest a much wider and dynamic venom landscape than initially appreciated in animals with a complex life cycle.

## Introduction

Venoms and the toxins they include are mostly used by animals for antagonistic interactions such as prey capture and defense from predators (Casewell et al., 2013; Fry et al., 2009). Pharmacological research is focused almost exclusively on the venoms of the adult stages of studied species despite the fact that many animals display remarkable transformations in body architectures and size along their development (Ruppert et al., 2004). Such vast differences can dictate very different interspecific interactions for different life stages (Wilbur, 1980). As venom is hypothesized to be metabolically expensive and in many cases highly specific (Casewell et al., 2013; Nisani et al., 2012), it is plausible that its composition might change between different life stages. Indeed, some ontogenetic variation was reported in the venoms of snakes (Gibbs et al., 2011) and cone snails (Safavi-Hemami et al., 2011), but up till now this phenomenon was not studied thoroughly in an animal with a complex life cycle throughout its development.

The oldest extant group of venomous animals is the marine phylum Cnidaria, which includes sea anemones, corals, jellyfish and hydroids. Cnidarians are typified by their stinging cell, the nematocyte that harbors a unique and highly complex organelle, the nematocyst (David et al., 2008; Kass-Simon and Scappaticci, 2002). This proteinaceous organelle is utilized as a miniature venom delivery platform (Lotan et al., 1995; Thomason, 1991). Most cnidarians have a complex life cycle that includes both sessile and mobile life stages of wide size distribution.

The starlet sea anemone *Nematostella vectensis* is becoming a leading cnidarian lab model as unlike many other cnidarian species it can be grown in the lab throughout its life cycle. This makes *Nematostella* a unique system to study the venom of an animal with a complex life cycle. Another advantage is that the high genetic homogeneity of the common *Nematostella* lab strain neutralizes individual genetic variation, which is far from trivial in most other venomous animals collected from the wild in limited numbers. Further, *Nematostella* has a sequenced genome and stage-specific transcriptomes and various molecular tools are available for its genetic manipulation (Helm et al., 2013; Layden et al., 2016; Putnam et al., 2007; Renfer et al., 2010; Wikramanayake et al., 2003). The *Nematostella* experimental toolbox is not unique only for cnidarians but also for venomous animals in general.

In the *Nematostella* life cycle, the female releases a gelatinous egg package and the male releases sperm to the water (Hand and Uhlinger, 1992). After the fertilization occurs, the zygote starts cleaving and forms in a few hours a blastula and less than 24 h post-fertilization (hpf) gastrulation is completed. A planula larva emerges from the egg package 48-72 hpf and starts swimming in the water. Six to seven days after fertilization the planula settles in soft substrate and starts to metamorphose into a primary polyp and sexual maturation takes about 4 months under lab conditions (Hand and Uhlinger, 1992) (Fig. 1A). Whereas the egg and planula are roughly spherical and measure only about 250 μm, the morphologically differentiated elongated adult polyp can reach the length of 4 cm in the wild and up to 20 cm in the lab (Hand and Uhlinger, 1992; Williams, 1975). Venom is produced in *Nematostella* by ectodermal gland cells and nematocytes (Moran et al., 2012b; Moran et al., 2013). Despite the fact that only at the sessile primary polyp stage *Nematostella* starts capturing prey and producing Nv1, a major sodium channel modulator toxin (Moran et al., 2012b; Moran et al., 2008a), nematocysts appear as early as 48 hpf in the swimming planula (Marlow et al., 2009). These are strong indications that venom might be present already in early life stages and that its composition might dramatically change across the *Nematostella* life cycle. Further complexity in the regulation of venom production might result from the fact that toxins in *Nematostella* are produced in different cell types that are unevenly distributed between various tissues (Moran et al., 2013) (Fig. 1B).

**Fig. 1.**
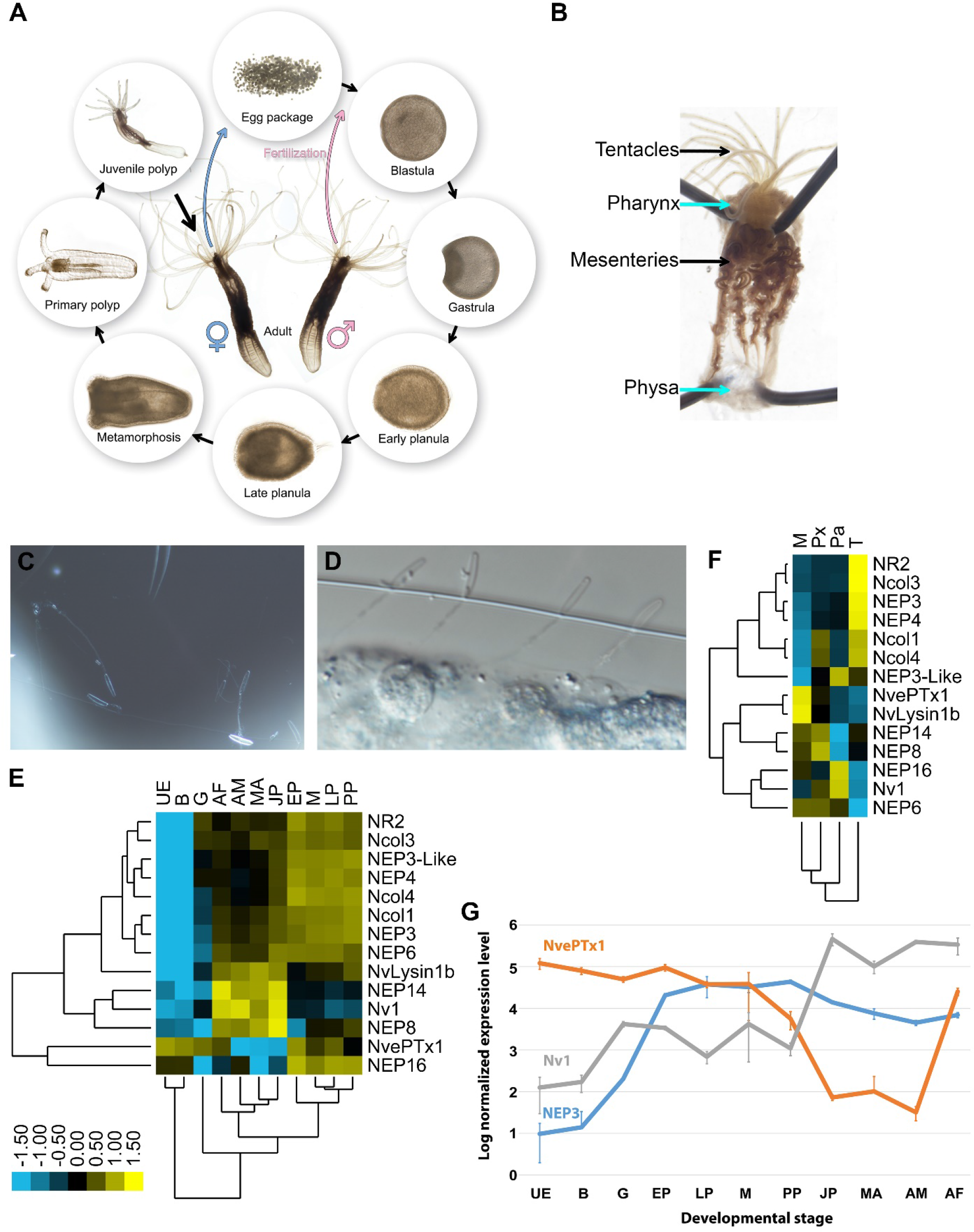
Expression of toxins across *Nematostella* development and tissues. (*A*) The life cycle of *N. vectensis*. (*B*) Dissected *Nematostella* female polyp. (*C*) Discharged planula nematocysts found in the medium after an encounter with A. salina nauplii. (*D*) Nematocysts of Nematostella planula pinned in the cuticle of an *A. salina* nauplii. (*E-F*) Heat maps of the nCounter differential expression levels of genes encoding toxins and other nematocyst proteins in various developmental stages and adult female tissues. (*G*) A graph at a logarithmic scale of the nCounter normalized expression levels of the genes encoding NEP3, NvePTx1 and Nv1 at each developmental stage. Each point is the average of three biological replicates and the error bars represent standard deviation. Key for panels E and G: UE=Unfertilized Egg; B=Blastula; G=Gastrula; EP=Early Planula; LP=Late Planula; M=metamorphosis; PP=Primary Polyp; JP=Juvenile Polyp; MA=Mixed Adults; AM=Adult Male; AF=Adult Female. Key for panel F: M=Mesenteries; Px=Pharynx; Pa=Physa; T= Tentacles.

Here, we carefully quantify and examine by multiple methods the spatiotemporal expression of known and novel *Nematostella* toxins across development and employ transgenesis to understand the dynamics of venom production in a species with a complex life cycle.

## Results

### Nematostella larvae are venomous

To assay the venomous potential of *Nematostella* larvae we incubated 4 days old swimming planulae with nauplii of the brine shrimp *Artemia salina.* Strikingly, within 10 min from the start of the incubation 3 out of 8 *Artemia* were paralyzed or dead, and within 90 min the vast majority (7 of 8) were dead (**Supplementary Movie S1**), whereas in a control group without planulae all *Artemia* were alive. This experiment revealed that *Nematostella* planulae are capable of rapidly killing a crustacean that is larger than they are. The relatively rapid effect and the size difference suggest that venom is involved in the process. Numerous discharged nematocysts were found in the water around the dead nauplii as well as in their cuticle (Figs. 1C-D), further suggesting that the stinging capsules are involved in the envenomation process.

### Distinct and dynamic expression patterns of toxin genes in Nematostella

To accurately measure the expression levels of known toxin genes, putative toxins and genes encoding nematocyst structural proteins we used the medium-throughput nCounter platform (see materials and methods), which was previously shown to exhibit high sensitivity and precision similar to that of real-time quantitative PCR (Prokopec et al., 2013). We assayed the RNA expression levels of the genes encoding the sodium channel modulator Nv1 (Moran et al., 2008a), the putative toxins NvePTx1, NEP3, NEP3-like, NEP4, NEP8 and NEP16 (Moran et al., 2013; Orts et al., 2013), the putative metallopeptidases NEP6 and NEP14 (Moran et al., 2013), the pore-former toxin NvLysin1b (Moran et al., 2012a) and the structural components of the nematocyst capsule Ncol1, Ncol3 and Ncol4 (David et al., 2008; Zenkert et al., 2011) as well as the putative nematocyst structural component NR2 (Moran et al., 2014). The RNA measurements were performed on nine developmental stages (Fig. 1E), adults of each sex and four dissected tissues of an adult female (Fig. 1F). The nCounter analysis revealed that many of the genes form informative clusters (Figs. 1E-F). It is noticeable that the expression patterns of NEP3, NEP3-like and NEP4 strongly clustered with those of genes encoding structural nematocyst components in both the developmental and tissue analyses (Figs 1E-F), which is consistent with the finding that these putative toxins are produced in nematocytes and are released from the capsule upon discharge (Moran et al., 2013). Other toxins such as Nv1, which was shown to be expressed in polyp ectodermal gland cells (Moran et al., 2012b), or Nvlysin1b, which was shown to be expressed in large gland cells in the pharynx and mesenteries since early age (Moran et al., 2012a), seemed to yield different expression patterns in the nCounter analysis and did not form large clusters (Figs. 1E-F). Comparing in detail the transcriptional expression of NEP3, Nv1 and NvePTx1, reveals strikingly distinct expression patterns across development (Fig. 1G). The expression of Nv1 is relatively low in early developmental stages and then sharply peaks in the juvenile and adult polyps to extraordinary levels that are higher by almost two orders of magnitude compared to the other toxins (Fig. 1G). These expression levels can be explained by the fact that the *Nematostella* genome contains more than a dozen gene copies encoding Nv1 (Moran et al., 2008b) and the peak in transcriptional expression late in development is consistent with earlier observations at the protein level (Moran et al., 2012b). In contrast to Nv1, NEP3 is expressed at high levels already from gastrulation, peaks in the early planula and remains roughly stable throughout the rest of development well into adulthood. Unlike the other toxins, the expression of NvePTx1 peaks at the unfertilized egg, drops sharply across development and rises again in the adult female (Fig. 1G).

### NvePTxl is a maternally deposited toxin

NvePTx1 was originally identified as a homolog of the type 5 potassium channel blocker BCsTx3 from the sea anemone *Bunodosoma caissarum* (Orts et al., 2013). We also identified bioinformatically several homologous sequences in the sea anemones *Anthopleura elegantissima* and *Metridium senile,* and the hydrozoan *Hydractinia symbiolongicarpus* (Fig. 2A). This suggests that this peptide family was already present in the last common ancestor of all Cnidaria but was lost multiple times in various cnidarian lineages.

**Fig. 2.**
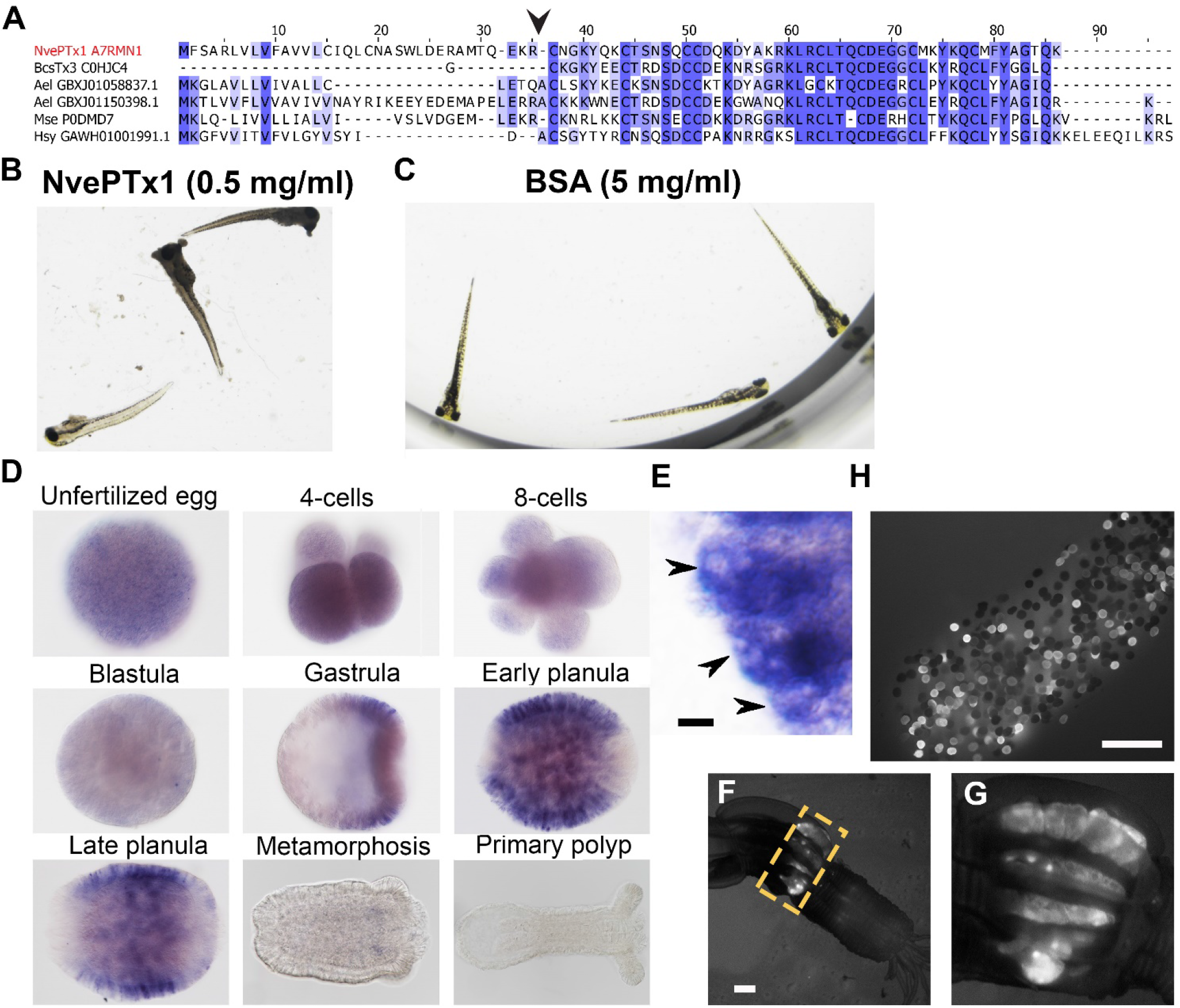
NvePTx1 is a toxin that is maternally deposited in *Nematostella* eggs. (*A*) Sequence alignment of NvePTx1 and its homologs from other cnidarian species. The arrowhead represents the proteolysis site that separates the mature toxin from the signal peptide. Accession numbers appear near each sequence. Nve=*Nematostella vectensis*; Bcs=*Bunodosoma caissarum*; Ael= *Anthopleura elegantissima*; Mse= *Metridium senile*; Hsy= *Hydractinia symbiolongicurpus*. (*B-C*) Zebrafish larvae 16 hours after incubation in NvePTx1 and BSA control. (*D*) Detection of *NvePTx1* expression by ISH. From the blastula stage on, the oral pole is to the right (*E*) Close up on planula stained by ISH for *NvePTx1* expression. Gland cells, which are discernible by their round vesicles that are free of stain, are indicated by arrows. Scale bar is 10μm (*F-G*) An adult F0 female polyp that as a zygote was injected by an *NvePTx1*::*mOrange2* construct. mOrange2 expression is noticeable in the gonads. Panel G is a close up of the region indicated in panel F by the yellow dotted box. Scale bar is 1000μm (*H*) An egg package laid by the female from panel F. Many of eggs express mOrange2. Scale bar is 1000μm.

First, to test if NvePTx1 is indeed a toxin we expressed it in a recombinant form and incubated zebrafish larvae with 0.5 mg/ml of highly pure recombinant peptide (assayed by reverse phase chromatography) for 16 hours. Upon the addition of the toxin, the fish reacted swiftly and started swimming extremely rapidly. Further, 10 fish larvae were dead by two hours from the beginning of the experiment and another 10 died within the next 18 hours (the end of the experiment) (Fig. 2B). The control group (incubated for 20 hours in 5 mg/ml bovine serum albumin) behaved normally throughout the experiment and all larvae were alive by its end (Fig. 2C). Next, to complement the nCounter experiment, we assayed the spatiotemporal expression pattern of NvePTx1 by in situ hybridization (ISH). We observed that while NvePTx1 is expressed uniformly throughout the unfertilized egg and early post-fertilization stages, in the gastrula the expression becomes spatially-localized and seems to be absent from the oral and aboral poles (Fig. 2D). In the planula, the expression is clearly observed in the ectoderm in packed gland cells absent from the two body poles, and upon metamorphosis the expression diminishes (Figs. 2D-E). The results of the ISH and nCounter experiments indicated that NvePTx1 is maternally deposited at the RNA level. Further, we could detect NvePTx1 peptide hits in supplementary datasets available from previous tandem mass spectrometry (MS/MS) analyses of *Nematostella* eggs (Levitan et al., 2015; Lotan et al., 2014), suggesting maternal deposition also at the protein level. To directly test this notion we have injected into *Nematostella* zygotes a transgenesis construct that carries the gene encoding the fluorescent reporter mOrange2 (Shaner et al., 2004) fused to an NvePTx1 signal peptide downstream of a putative NvePTx1 promoter. Noticeably, several females of the injected first generation (F0) exhibited strong expression of mOrange2 in round structures in their mesenteries, which are most probably the ovaries where the eggs are formed (Fig. 2F-G). This observation is congruent with the fact that the mesentery is the only female adult tissue where high NvePTx1 transcript levels are detected at high levels by the nCounter analysis (Fig. 1F). Further, upon induction of spawning, the female polyps with the fluorescent mesenterial tissue released egg packages harboring plenty of fluorescent eggs (Fig. 2H), strongly supporting maternal deposition of NvePTx1.

### Phylogeny, primary structure and activity of NEP3

The NEP3, NEP4, and NEP8 gene products were previously detected by MS/MS to be released from *Nematostella* nematocysts upon discharge and hence were hypothesized to be putative toxins (Moran et al., 2013). We detected an additional gene encoding a NEP3 homolog in the *Nematostella* genome and named it NEP3-like. The four nucleotide sequences were translated *in silico* and were suggested to encode precursors of secretory proteins equipped with typical signal peptides. Search in the Pfam database (Finn et al., 2010) showed that each precursor contains three ShKT sequence motives (PF01549, **Fig 3A**) typical for several potent cnidarian toxins (Aneiros et al., 1993; Castaneda and Harvey, 2009). Based on these common features and their sequence similarity, we designated NEP3, NEP3-like, NEP4, and NEP8 as the “NEP3 family”. Search in publicly available transcriptomic shotgun assembly databases led to identification of several sequences of homologs from the sea anemones *Edwardsiella lineata, Aiptasia pallida* and *Anthopleura elegantissima* as well as the stony corals *Acropora digitifera* and *Stylophora pistillata,* showing significant similarity and identical domain structure to the NEP3 family members from *Nematostella* (**Supplementary Fig. S1**). A phylogenetic analysis revealed that each of NEP3, NEP4 and NEP8 from *Nematostella* form a strongly supported (bootstrap values >0.5) clade with a highly similar protein from *Edwardsiella* (Fig. 3B), indicating orthology. Thus, the new sequences from *E. lineata* were named according to their *Nematostella* orthologs. As *Nematostella* and *Edwardsiella* are members of the basally-branching sea anemone family Edwardsiidae (Rodriguez et al., 2014; Stefanik et al., 2014) and they both possess NEP3, NEP4 and NEP8 we can infer that those three proteins probably originated in the last common ancestor of the Edwardsiidae. Other cnidarian species bear more distantly related NEP3 family members, and their exact orthologous or paralogous nature could not be determined due to low bootstrap values (Fig. 3B). However, their presence in stony corals suggests that those proteins already appeared 500 million years ago in the last common ancestor of stony corals and sea anemones (Shinzato et al., 2011), but were lost in multiple hexacorallian lineages.

**Fig. 3.**
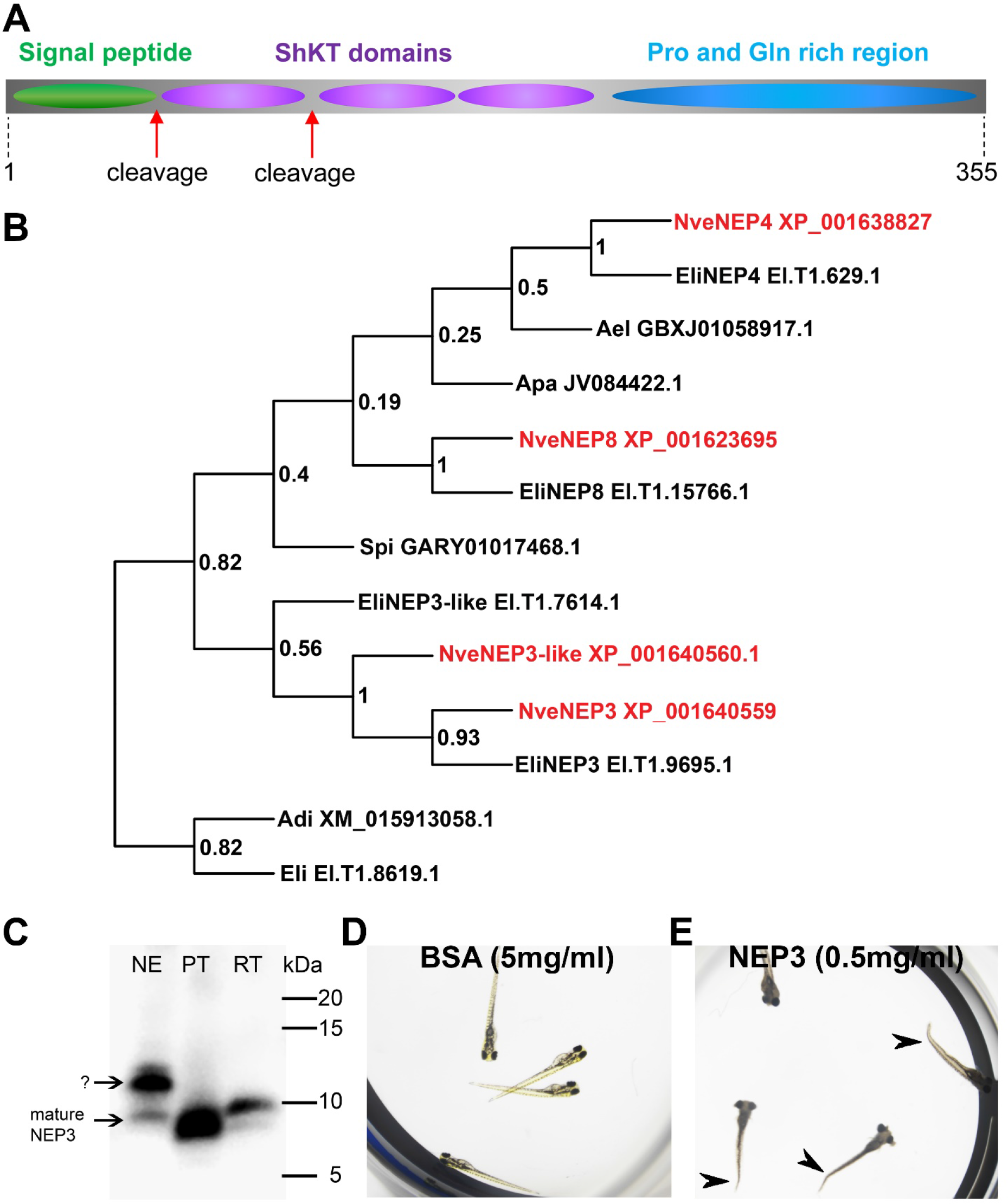
NEP3 is a toxin that is processed from a long precursor protein and is part of a large family. (*A*) The primary structure of the NEP3 precursor. (*B*) A maximum-likelihood tree of the NEP3 family. Accession numbers appear near each sequence and bootstrap values (fraction of 1000 bootstraps) appear near each node. Nve=*Nematostella vectensis*; Eli:*Edwardsiella lineata*; Ael=*Anthopleura elegans*; Apa=*Aiptasia pallida*; Spi=*Stylophora pistillata*; Adi=*Acropora digitifera*. (*C*) Western blot with rat Anti-NEP3 antibody. Samples are discharged nematocyst extract (NE), native purified toxin (PT) and recombinant toxin (RT). (*D-E*) Zebrafish larvae after 16 hours incubation in NEP3 and BSA control. Arrowheads are pointing at twitched tails.

To check whether NEP3 serves as a toxin we planned to express it in a recombinant form. However, it was initially not clear in what native form this protein is found in the animal as we detected a potential Lys-Arg tandem, which is a prominent cleavage signal in nematocyst proteins (Anderluh et al., 2000), between the first and second domains of NEP3. Hence, we decided to first explore the primary structure of the native mature NEP3. For this aim, we discharged lyophilized nematocysts and analyzed the molecular weight of NEP3 making part of the ejected protein mixture by western blot with custom polyclonal antibodies against the first NEP3 ShKT domain. However, the western blot resulted in two major bands (∼10 and 12 kDa) (Fig. 3C). A gradual three-step FPLC procedure of gel filtration, anion exchange and reverse phase chromatography was applied to purify the dominant NEP3 fragment, and at each stage we carried on with the fraction with the strongest western blot signal. This procedure yielded a nearly pure fraction of mature NEP3 fragment with molecular weight ∼10 kDa. Comparison of the electrophoretic mobility of the native mature NEP3 and a recombinant peptide corresponding to the first domain showed that the two peptides possess highly similar molecular weights (Fig. 3C). Thus, we conclude that a native peptide composed of only the first domain of NEP3 is found in *Nematostella* and is released from the nematocyst upon discharge. Following this finding we carried out a toxicity test where we incubated zebrafish larvae with 0.5 mg/ml recombinant mature NEP3. While in the control group (incubated for 20 hours in 5 mg/ml bovine serum albumin) all the 20 fish survived (Fig. 3D), all 17 fish larvae in the NEP3-treated group died within 5 hours and the larvae exhibited pronounced contraction and tail twitching (Fig. 3E) that might suggest the mature NEP3 peptide is neurotoxic.

### Different nematocytes express different NEP3 family members

To gain improved resolution of the expression of the four NEP3 family members we employed ISH to localize their expression in five developmental stages. All the four genes are expressed in nematocytes on the surface of the early planula (Fig. 4A; Supplementary Fig. S2). However, beginning at the late planula stage the expression of NEP8 shifts to only a handful of nematocytes in the lower pharynx (Fig. 4A). In contrast to NEP8, the three other family members, NEP3, NEP3-like and NEP4 are expressed throughout the ectoderm in nematocytes, with high concentration of expressing cells in the oral pole. In the primary polyp, expression of NEP3, NEP3-like and NEP4 is noticeable in nematocytes in the body wall and physa ectoderm and in the upper and lower pharynx (Fig. 4A). Further, in tentacle tips, which are very rich with nematocytes, there are large numbers of nematocytes expressing the three toxins, fitting well our nCounter results (Fig. 1F). A superficial look on these results might give the impression that in *Nematostella* there is one small population of pharyngeal nematocytes that express NEP8 and another very large population of nematocytes that express NEP3, NEP3-like and NEP4 in much of the ectoderm. However, we decided to use double fluorescent ISH (dFISH) to check if this is truly the case.

**Fig. 4.**
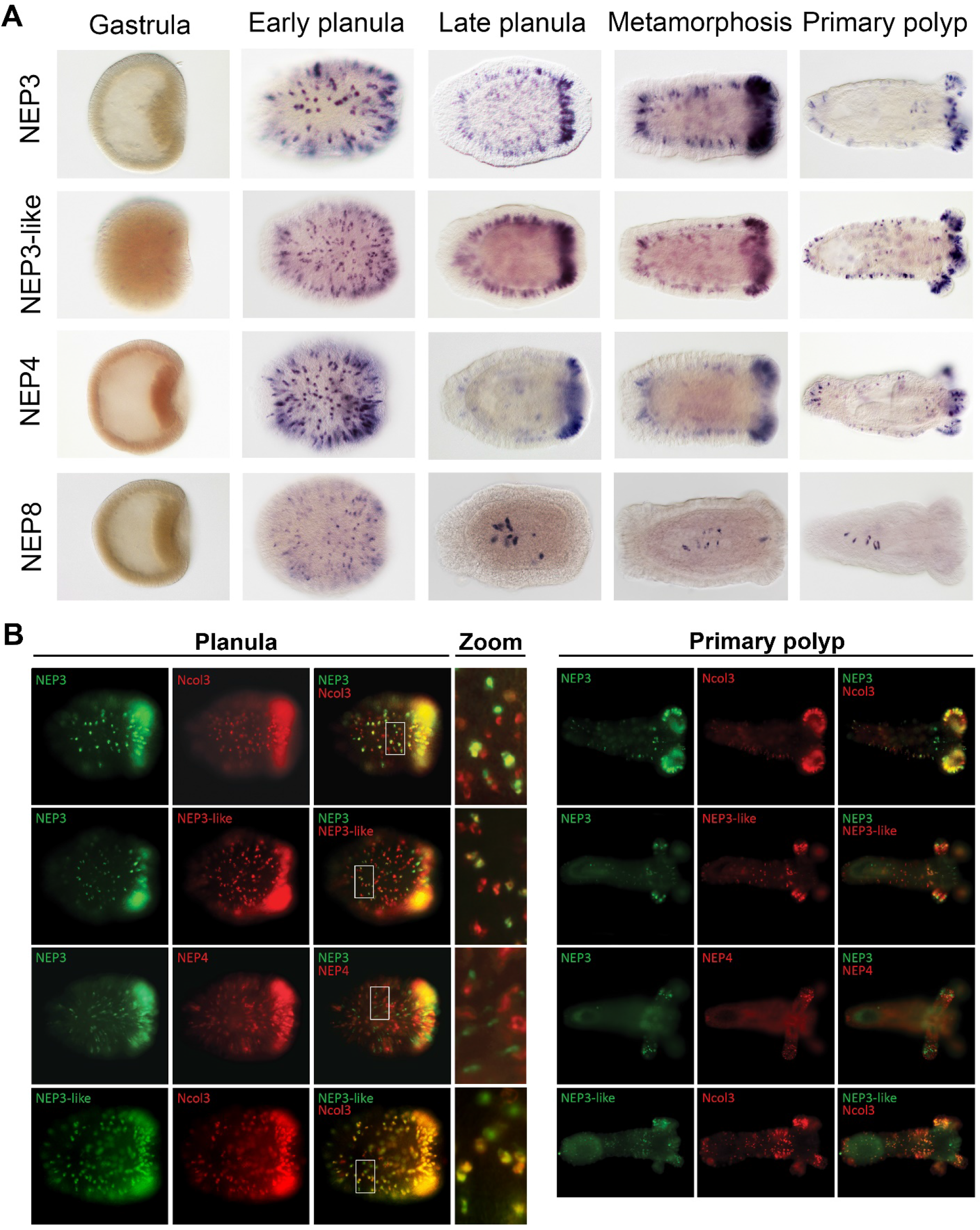
Partially overlapping and distinct expression patterns of NEP3 family members. (*A*) Expression of the NEP3 family members in five developmental stages of *Nematostella* as determined by ISH. (*B*) Expression of the NEP3 family members in planulae and polyps of Nematostella as determined by dFISH. In all panels, the oral pole is to the right.

In the dFISH experiments we localized the mRNA combinations of NEP3 with Ncol3, NEP3 with NEP3-like, NEP3 with NEP4, and NEP3-like with Ncol3. Unexpectedly, all the combinations showed only limited overlap in their expression, with NEP3-like and Ncol3 showing the highest overlap and NEP3 and NEP4 showing the lowest (Fig. 4B). This result can be explained by two non-exclusive explanations: the first is that the nematogenesis (production nematocysts) is a highly dynamic process that requires different genes to be expressed at different times along the nematocyst maturation process; the second is that the three family members are mostly expressed in different nematocyte populations and only few nematocytes express all family members.

To test the latter hypothesis we injected *Nematostella* zygotes with a construct carrying a chimera of the signal peptide of NEP3 with mOrange2 downstream of the putative promoter of NEP3. Zygotes started expressing mOrange2 in nematocytes about 4 days after injection. We raised the positive F0 animals to adulthood and then crossed them with wildtype polyps to generate founders. We found 6 female and 3 male founders that the mOrange2 expression in their F1 progeny imitated the expression of the native gene (Fig. 5A-C). To verify this we performed double ISH on F1 as well as F2 animals (third generation) and found nearly perfect overlap between the transcriptional expression of the mOrange2 transgene and that of the NEP3 toxin gene (Fig. 5D-F). mOrange2 expression was observed in many nematocytes with especially dense populations in the tentacle tips and the mesenteries (Fig. 5C). Strong expression of mOrange2 was also detected in numerous nematocytes within the nematosomes (Fig. 5G-I), defensive structures that *Nematostella* releases to its surroundings and within egg packages (Babonis et al., 2016).

**Fig. 5.**
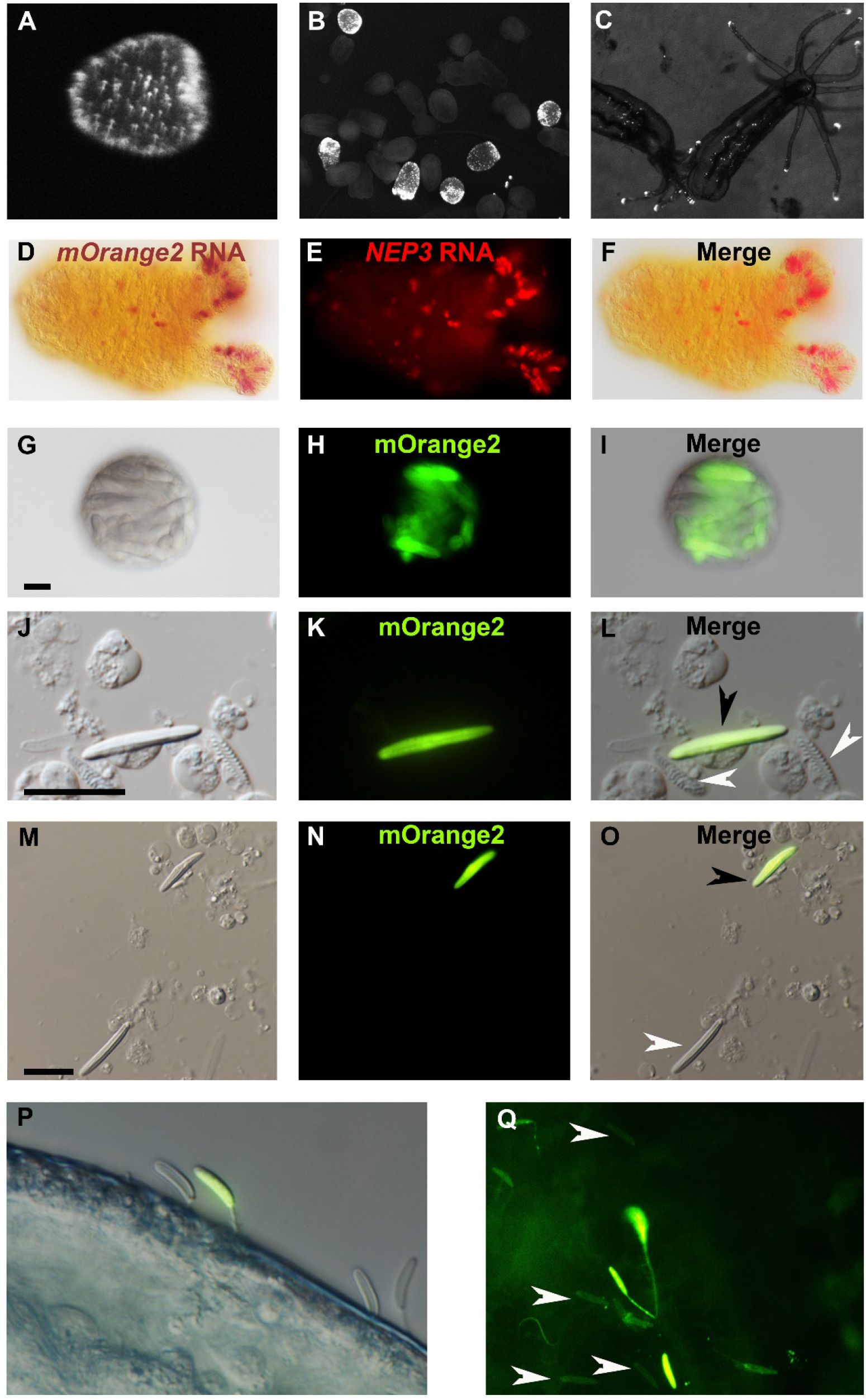
NEP3 is expressed only in a subset of nematocytes. (*A*) 3 days of planula of an F1 transgenic line expressing mOrange2 under a *NEP3* promoter. (*B*) 6 days old metamorphosing larva of the same transgenic line. (*C*) 2 months old juvenile polyps of the same line. (*D-F*) Double ISH of an F2 primary polyp of the transgenic line with probes against NEP3 and mOrange2. mOrange2 transcripts were stained with an NBT (*nitro-blue tetrazolium chloride*) and BCIP (5-bromo-4-chloro-3’-indolyphosphate p-toluidine) solution that forms purple crystals, whereas NEP3 transcripts were stained with FastRed that provides fluorescent red signal. (*G-I*) A nematosome of the transgenic line exhibiting mOrange2 fluorescent signal within its nematocytes. (*J-L*) A spread of dissociated cells of the tentacles of an F2 transgenic NEP3 polyp. An mOrange2-positive nematocyte is indicated by a black arrow and negative spirocytes are indicated by white arrows. (*M-O*) A picture of another field of the cell spread from previous panels. An mOrange2-positive nematocyte is indicated by a black arrow and a negative nematocyte is indicated by a white arrow. Scale bar is 10μm in panels G-O. (*P*) Nematocytes of a transgenic F2 polyp pinned in the cuticle of *A. salina*. Only one nematocyte in the picture is mOrange2-positive. (*Q*) Nematocytes of an F2 polyp of the same transgenic line are pinned in the skin of a zebrafish larva. Only some of them are mOrange2 positive. White arrows indicate examples for negative nematocytes.

Next, we dissociated tentacles of transgenic F1 animals by a mixture of commercial proteases into single cells and observed what cells express mOrange2 and hence NEP3. As expected, we observed that NEP3 is expressed in nematocytes, but not in spirocytes (Fig. 5J-L), which are believed to be utilized for adherence and not for venom delivery (Mariscal et al., 1977). Moreover, we also observed that NEP3 is expressed only in a subpopulation of nematocytes (Fig. 5M-O), suggesting like the ISH and dFISH experiments that different nematocytes express different toxins.

However, at that point there was a possibility that the mOrange2 is noticeable only in developing nematocytes and that mature capsules are not glowing due to various technical limitations such as the mature capsule wall obstructing light. In order to test whether there are mature mOrange2-positive capsules in our transgenic line we challenged the F1 polyps with *Artemia* nauplii and zebrafish larvae. We then took the attacked prey items and visualized them with fluorescent microscopy. Strikingly, mOrange2-positive capsules with glowing tubules were pinned in the crustacean and fish cuticle or skin, respectively (Fig. 5P-Q), proving that those are mature capsules. However, these capsules were accompanied by other capsules that were mOrange2-negative, strongly suggesting once again that only a certain nematocyte subpopulation in *Nematostella* is expressing NEP3.

## Discussion

Our finding of vastly different expression levels of toxins in different developmental stages and adult tissues, strongly suggests that venom composition changes across development and that each arsenal of toxins might have been shaped by selection for different biological needs. As *Nematostella* develops from a non-predatory, swimming planula larva to an adult predatory polyp that is 150-fold larger than the larva (Fig. 1A), its interspecific interactions vastly change across development. We hypothesize that these dynamic interactions, coupled with the potentially high metabolic cost of toxins (Nisani et al., 2012), have driven the evolution of a distinct venom composition in each developmental stage. Moreover, in the case of cnidarians, an additional metabolic cost stems from the fact that nematocysts are single-use venom delivery apparatuses that have to be reproduced in very high numbers after each antagonistic interaction. Hence, there is a clear advantage in using a highly adapted venom in each developmental stage.

Because venom in planula is used purely for defense, whereas the venom in the polyp is used both for defense and for prey capture, our results also relate to previous results in scorpions (Inceoglu et al., 2003) and cone snails (Dutertre et al., 2014) where venom compositions for defense and prey-capture were shown to differ. Further, some of the toxins we localized can be attributed for specific functions based on their expression patterns. For example, it is clear the NvePTx1 is a defensive toxin due to its occurrence in the egg and in ectodermal gland cells of the planula (Figs. 2D-H and 6), whereas NEP8 is probably used in the polyp for killing prey after it is swallowed, as it is expressed exclusively in the lower pharynx and mesenteries (Figs. 4A and 6).

**Fig. 6.**
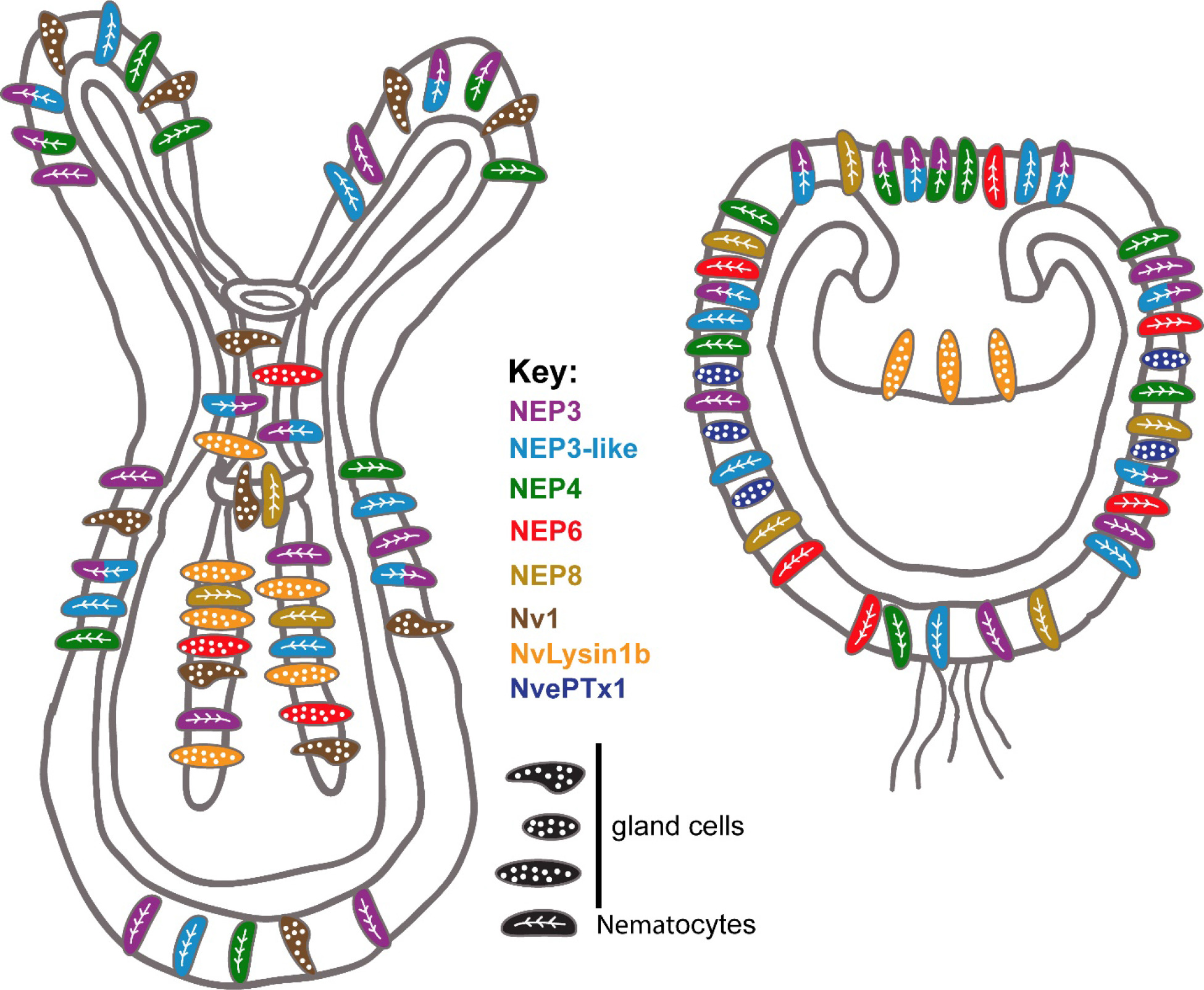
Summary of the current knowledge on spatiotemporal toxin expression in early planula and primary polyp of *Nematostella*. This illustration summarizes the current work as well as previous works (Moran et al., 2012a; Moran et al., 2012b; Moran et al., 2013).

Chemical protection of the eggs was reported in several animals such as black widow spiders (Buffkin et al., 1971; Lei et al., 2015), snails (Dreon et al., 2013), octopuses and some fishes and amphibians (Bane et al., 2014). Our data on *Nematostella* NvePTx1 expressed in eggs and embryonic stages further support the idea of ecological importance of chemical defence in early life stages across the animal kingdom. Our findings that NEP3 family members are expressed in different population of nematocytes reveals that nematocyte diversity in *Nematostella* exceeds well-beyond the morphology-based assessments, which revealed only two types of nematocysts in *Nematostella:* basitrichous haplonemas (also called basitrichs) and microbasic mastigophores (Frank and Bleakney, 1976; Zenkert et al., 2011). A study in Hydra discovered that two members of a single pore-forming toxin family are expressed in two morphologically distinct types of nematocytes (Hwang et al., 2007), but to the best of our knowledge, similar complexity of toxin expression patterns in morphologically similar nematocytes was never reported before. Further, mechanisms of venom biosynthesis at cellular level resolution have not been reported in more complex venomous organisms as well. However, at lower resolution, some reports suggest that within one venom gland different secretory units are specialized on production of a limited number of toxins (Dutertre et al., 2014; Undheim et al., 2015). Thus, specialized venom secretory cells is probably a common trait among venomous animals.

The distinct expression patterns of the NEP3 family also provides important indications regarding toxin evolution. For example, the expression of NEP8 in pharyngeal nematocytes, and its absence from the tentacles and outer body wall, where its paralogs NEP3, NEP3-like and NEP4 are expressed (Figs. 4A and 6), is an indication for sub-or neo-functionalization. The specialization of the different family members is also supported by their conservation in *Edwardsiella* (Fig. 3B).

Variation in expression patterns of the NEP3 family members and the fact that at least four different types of gland cells at distinct developmental stages and tissues express different toxins (Nv1, Nvlysin1b, NEP6 and NvePTx1) in *Nematostella* suggests a highly complex venom landscape in this species (Fig. 6). At first glance, such a system might seem relatively inefficient. However, we hypothesize that harboring many different toxin-producing cell types, provides modularity and enables evolutionary plasticity of toxin expression. Indeed, our results as well as results of others, suggest that different sea anemones species express similar toxins in different cell types (Moran et al., 2012b) and different tissues (Macrander et al., 2016). This evolutionary plasticity might be one of the factors that made sea anemones such a successful group that inhabits all the world’s oceans for the last 600 million years.

## Materials and Methods

### Animals

*Nematostella* embryos, larvae and juveniles were grown in 16%o sea salt water at 22°C. Adults were grown in the same salinity but at 17°C. Polyps were fed with *A. salina* nauplii three times a week. Induction of gamete spawning was performed according to a published protocol (Genikhovich and Technau, 2009b).

### nCounter analysis

Total RNA from different developmental stages and body parts of adult female *Nematostella* was extracted with Tri-Reagent (Sigma-Aldrich, St. Louis, MO, USA) according to manufacturer’s protocol, treated with Turbo DNAse (Thermo Fisher Scientific, Waltham, MA, USA) and then re-extracted with Tri-Reagent. RNA quality was assessed on Bioanalyzer Nanochip (Agilent, Santa Clara, CA, USA) and only samples with RNA Integrity Number (RIN) ≥ 8.0 were used. Those samples were analyzed on the nCounter platform (NanoString Technologies, Seattle, WA, USA; performed by Agentek Ltd., Israel) in triplicates as previously described (Geiss et al., 2008). In brief, for each transcript to be tested, two probes were generated and hybridized to the respective mRNA. The mRNAs were immobilized on a cartridge and the barcodes on one of the probes were counted by an automated fluorescent microscope. For normalization we used a geometric mean of the expression levels of 5 reference genes with stable expression across development. The genes were selected as follows: we calculated the Shannon entropy (as described in (Schug et al., 2005)) for each of 23,041 *Nematostella* genes based on normalized transcript abundance estimates for six time-points of *Nematostella* development (Helm et al., 2013). We then ranked the genes by entropy, which indicates minimal temporal change in abundance, and from the top 20 chose five genes (NCBI Reference Sequences XM_001629766.1, XM_001628650.1, XM_001625670.1, XM_001640487.1 and XM_001624235.1) with complete gene models and mean abundance levels spanning the expected experimental range. Probe sequences, entropy scores and all raw and normalized nCounter read data are available in Supplementary Table S1.

### In situ hybridization (ISH)

Single and double ISH were performed as previously described (Genikhovich and Technau, 2009a; Moran et al., 2013). dFISH was performed also according to published protocols (Nakanishi et al., 2012; Wolenski et al., 2013) with tyramide conjugated to Dylight 488 and Dylight 594 fluorescent dyes (Thermo Fisher Scientific). In ISH and FISH, embryos older than 4 days were treated with 2u/μl T1 RNAse (Thermo Fisher Scientific) after probe washing in order to reduce background. Stained embryos and larvae were visualized with an Eclipse Ni-U microscope equipped with a DS-Ri2 camera and an Elements BR software (Nikon, Tokyo, Japan).

### Transgenesis

To generate transgenic constructs we replaced the mCherry gene with mOrange2 (Shaner et al., 2004) and replaced the promoter sequence of the pNvT-MHC::mCH plasmid (Renfer et al., 2010). For the *NEP3* gene, we inserted to the plasmid 920 bp upstream of the transcription start site as well as the non-coding first exon, first intron and the part of the second exon that encodes the signal peptide of NEP3 (scaffold_7:1,219,288-1,221,320 of the *Nematostella* genome). For the NvePTX1 gene, we inserted to the plasmid 1033 bp upstream of the transcription start site as well as the non-coding first exon, first intron and the region of the second exon that encodes the signal peptide (scaffold_14:1,246,079-1,247,853 of the *Nematostella* genome). The constructs were injected with the yeast meganuclease *I-SceI* (New England Biolabs, Ipswich, MA, USA) to facilitate genomic integration (Renfer et al., 2010). Transgenic animals were visualized under an SMZ18 stereomicroscope equipped with a DS-Qi2 camera (Nikon).

### Phylogenetics

Sequences of NEP and NvePTx1 protein families were retrieved using blast searches (Altschul et al., 1990) against NCBI’s non-redundant nucleotide sequence database and the EdwardsiellaBase (Stefanik et al., 2014). Maximum-likelihood analysis was employed for the reconstruction of the molecular evolutionary histories. Trees were generated using PhyML 3.0 (Guindon et al., 2010), and node support was evaluated with 1,000 bootstrapping replicates.

### Tentacle Dissociation

Tentacles of *Nematostella* were dissociated using a combination of papain (3.75 mg/ml; Sigma-Aldrich: P4762), collagenase (1000 U/ml; Sigma-Aldrich: C9407) and pronase (1 mg/ml; Sigma-Aldrich: P5147) in DTT (0.1 M) and PBS solution (10 mM sodium phosphate, 150 mM NaCl, pH 7.4). The tentacles were incubated with the protease mixture at 22°C overnight. The tissues were then dissociated into single cells by flicking the tubes gently and then by centrifugation at 400 × *g* for 15 minutes at 4°C, followed by resuspension in PBS.

### Purification of NEP3 from nematocysts

Lyophilized nematocysts were obtained from Monterey Bay Labs (Caesarea, Israel). 2.5 g of the nematocysts were discharged by incubation with 80 ml of 1% sodium triphosphate (Sigma-Aldrich, USA). Following centrifugation (21,000 × g, 20 min), the crude extract was concentrated with Amicon centrifugal filters with 3 kDa cut off (Merck Millipore, Billerica, MA, USA) to 2 ml volume, filtered through Amicon centrifugal filters with 50 kDa cut off (Merck Millipore) and used for further purification. At the first step, the extract was fractionated by size exclusion FPLC on a calibrated Superdex 75 column (60×1.6 cm, GE Healthcare, Little Chalfont, UK) in PBS buffer. Protein fractions with molecular weight less than 17.6 kDa were pooled and the PBS buffer was exchanged to 20 mM ethanolamine pH 9 using Amicone centrifugal filters, cut off 3 kDa. At the second step, the SEC fractions were separated by anion exchange FPLC using a HiTrapQ HP column (1 ml, GE Healthcare) and a NaCl concentration gradient (0-750 mM NaCl in 30 column volumes, 20 mM ethanolamine pH 9.0). Fractions were analysed by western blot with anti-Nep3 antibodies and positive ones were pooled. At the last step, Nep3 fragment was purified by reverse phase FPLC on a Resource RPC column (3 ml, GE Healthcare) using acetonitrile concentration gradient (8-60% CH3CN in 25 column volumes, 0.1% trifluoracetic acid). Fractions corresponding to individual peaks were collected and analysed by western blot with anti-NEP3 antibodies.

### Recombinant expression and purification of toxins

A synthetic DNA fragment encoding the full NEP3 polypeptide was purchased from GeneArt (Regensburg, Germany). The fragment corresponding to the first domain of NEP3 between the Lys-Arg cleavage sites was amplified by PCR, cloned and expressed as a His6-thioredoxin fusion protein in Shuffle T7 *Escherichia coli* strain (New England Biolabs).

NvePTx1 synthetic DNA fragment was purchased from Integrated DNA Technologies (Coralville, IA, USA) and cloned into a modified pET40 vector (fragment encoding DSBC signal peptide was erased from it by Protein Expression and Purification facility of the Hebrew University to allow cytoplasmic expression of DSBC). NvePTx1 was expressed in BL21(DE3) *E. coli* (Merck Millipore) strain as a fusion with His6-DSBC.

The polyhistidine tag of the fusion proteins was used for purification from the *E. coli* lysate by nickel affinity FPLC. Purified fusion proteins were cleaved into two fragments by Tobacco Etch Virus (TEV) protease (room temperature, overnight) at a TEV protease cleavage site upstream the toxin fragments. The recombinant toxins were then purified by reverse phase FPLC on a Resource RPC column (GE Healthcare) using an acetonitrile concentration gradient in 0.1% trifluoroacetic acid.

### Western blot

Custom polyclonal antibodies specific to the first domain of NEP3 were purchased from GenScript (Piscataway, NJ, USA). Synthetic peptide containing the amino acid positions 47-91 was used as an antigen for immunization of two rats. The antibodies were affinity purified of a column coated with the antigen. Proteins were separated by electrophoresis on 10-20% gradient Tris-tricine gels (Bio-Rad, Hercules, CA, USA) and consequently transferred to 0.2 um PVDF membranes (Bio-Rad). Membranes were blocked by 5% skim milk in TBST buffer (50 mM Tris base, 150 mM NaCl, 0.1% Tween 20, pH 7.6) and incubated with anti-NEP3 antibodies (1 ug/ml) in 5% Bovine serum albumin (BSA) in TBST buffer (4^0^C, overnight). This was followed by incubation with goat anti-rat IgG antibodies conjugated with horseradish peroxidase (0.1 ug/ml; Jackson ImmunoResearch, West Grove, PA, USA) in 5% skim milk in TBST (room temperature, overnight). ECL reagent (GE Healthcare) was used for visualization of the protein bands interacting with anti-NEP3 antibodies. Chemiluminescence was recorded with an Odyssey Fc imaging system (LI-COR Biosciences, Lincoln, NE, USA) and fluorescent size marker (Bio-Rad) was imaged on the same system.

### Toxicity assays

To assess the toxicity of putative toxins, 4 days old *Danio rerio* larvae (generously provided by Dr. Adi Inbal, the Hebrew University medical school) were incubated with 0.5 mg/ml peptides in 500 μl well. Each experiment was conducted in duplicates. 5 mg/ml BSA was used as a negative control. Effect was filmed and monitored under an SMZ18 (Nikon) stereomicroscope after 5 min, 15 min, 1 h, and 15-17 h of incubation.

## Author Contributions

YYCS, MYS, KS and YM designed the research. YYCS, MYS, AF, VM and KS performed the research; YM analyzed the data; MYS, YYSC and YM wrote the paper.

## Acknowledgements

We are grateful to Dr. David Fredman (Department of Informatics, University of Bergen) for his invaluable help with quantitative and computational methods. We are also grateful to Dr. Mario Lebendiker and Dr. Tsafi Danieli (Protein Expression and Purification Facilities of the Hebrew University) for their help with recombinant expression and chromatography and for the help of Dr. Dana Reichmann and Dr. Bill Breuer (Department of Biological Chemistry of the Hebrew University) for their help with mass spectrometry. K.S. was supported by a Marie Sklodowska-Curie Individual Fellowship (654294). This work was supported by Israel Science Foundation grant no. 691/14 and Binational Science Foundation grant no. 2013119 to Y.M.

## Supporting information

**Supplemental movie 1** (WMV format). The interaction between *Nematostella* planulae and *Artemia* nauplii. This movie is related to Fig. 1

**Supplementary Table S1** (XLS format). Information of nCounter probe sequences, nCounter raw and normalized data and entropy list for *Nematostella* transcripts. This table is related to Fig. 1.E-G.

**Supplementary Fig. S1** (PDF format). Multiple sequence alignment of the NEP3 family members from Fig. 3B. Accession numbers appear near each sequence. *Nematostella* sequences are in red.

**Supplementary Fig. S2** (PDF Format). Close up on ISH staining of NEP3, NEP3-like, NEP4 and NEP8 reveals that they are expressed in nematocytes. Scale bars are 20μm. This figure is related to Fig. 4.

